# One Hundred Years of Influenza A Evolution

**DOI:** 10.1101/2024.02.27.582392

**Authors:** Bjarke Frost Nielsen, Christian Berrig, Bryan T. Grenfell, Viggo Andreasen

## Abstract

Leveraging the simplicity of raw nucleotide distances, we provide an intuitive window into the evolution of the human influenza A ‘nonstructural’ (NS) gene. In an analysis suggested by the eminent Danish biologist Freddy B. Christiansen, we illustrate the existence of a continuous genetic “backbone” of influenza A NS genes, steadily increasing in distance to the 1918 root over more than a century. Interestingly, the 2009 influenza pandemic represents a clear departure from this enduring genetic backbone. Utilizing nucleotide distance maps and phylogenetic analyses, we illustrate remaining uncertainties regarding the origin of the 2009 pandemic, highlighting the complexity of influenza evolution. The NS gene is interesting precisely because it experiences neutral genetic drift over long periods of time time, while sudden deviations from this drift pattern can indicate changes in other genes via the hitchhiking effect. Our approach employs two measures based on genotypic distance — the rooted temporal Hamming map and the unrooted temporal Hamming distribution — to analyze the evolutionary dynamics of the NS gene. The rooted Hamming map elucidates distances between a reference sequence and all other sequences over time. In contrast, the unrooted temporal Hamming distribution captures the distribution of genotypic distances between simultaneously circulating viruses, thereby revealing patterns of sequence diversity and epi-evolutionary dynamics. Our study aims to supplement traditional tree-based phylogenetic inference with these direct temporal distance-based measures, offering transparent insights into the evolution of the influenza NS gene.

## I. INTRODUCTION

In the late 1990s and early 2000s, genetic material originating from the influenza strain which caused the severe 1918 ‘Spanish flu’ pandemic was isolated at Brevig Mission, Alaska (previously known as Teller Mission).The individual gene sequences were subsequently published and analyzed [1–8]. While the 1918 H1N1 pandemic hit the Brevig Mission settlement hard, killing the majority of the inhabitants [9], such extreme fatality rates were not uncommon in Alaskan communities struck by the 1918 influenza [10–12]. The high level of preservation by the Alaskan permafrost was what allowed modern sequencing of 1918 influenza genes, including the NS gene which is our main object of study. More broadly, the isolation and genetic characterization of this historic pathogen opened up new windows into the evolution and pathogenicity of influenza A [13–18].

In 1986, a decade before the isolation of the 1918 strain, Buonagurio et al. [19] had employed a nucleotide distance metric to study the long-term evolution of influenza A over an approximately 50 year period (1933-1985), viewed through the NS (*‘nonstructural’*) gene. In this study, we extend the analysis of the evolution of the NS gene to more than a century (1918-2023), using two complementary nucleotide distance metrics. The current study is motivated by an analysis that Freddy B. Christiansen carried out in 2001-2002 in connection with our work on the interaction between influenza drift and herd immunity [20], which had prompted us to study the work of Buonagurio et al. [19]. At that time, Freddy had moved to the new Bioinformatics Research Center (BIRC) at Aarhus University to get in closer contact with the molecular aspects of population genetics and was familiarizing himself with hands-on projects of sequence data-analysis. When the sequence of the NS gene from the Brevig Mission virus was published [4], Freddy immediately saw the possibility to extend Buonagurio’s time line back another couple of decades.

### Influenza classification

Before we continue to the main analysis, we will briefly introduce the nomenclature and classification of influenza A viruses, and comment on the usefulness of focusing on the NS gene, rather than the more prominent HA and NA genes.

All influenzaviruses are members of the *Orthomyxoviridae* family, making up the *Alpha-, Beta-, Gamma-* and *Deltainfluenzavirus* genera. Each genus has just a single member, the Influenza A, B, C and D viruses, respectively. Of these, influenza A is the most diverse and the one responsible for all known human influenza pandemics [21].

The genome of influenza A is segmented and consists of 8 strands (or genes), two of which – HA and NA – code for significant antigens – hemagglutinin and neuraminidase. Although our focus will be on the NS gene, we first discuss the role of HA and NA since these two genes and their corresponding antigens largely determine the natural selection induced by herd immunity and form the basis for subtype classification. Since the level of population immunity varies dramatically over time, HA and NA evolution shape the evolutionary background in which the role of the NS gene must be interpreted.

Influenza A *subtypes* are antigenically characterized by their surface proteins (H/HA) and (N/NA) leading to designations such as H3N2 (a subtype displaying a type-3 hemagglutinin and a type-2 neuraminidase protein). As of mid-2023, 18 hemagglutinin (H) and 11 neuraminidase (N) subtypes have been identified [22].

The significance of the different Influenza A subtypes is immunological: different subtypes show limited cross-reaction in serological tests, and subtypes with distinct HA molecules do not cross-react. As such, the risk of a large outbreak or pandemic is greatly increased by the emergence of a subtype whose HA molecule is not related to those of influenza viruses in widespread circulation immediately before it [23].

### Antigenic shift and drift

When the new subtypes H1N1, H2N2 and H3N2 appearred in 1918, 1957 and 1968, the previous subtype disappeared during the associated pandemic – although the underlying mechanism is poorly understood [24, 25]. These events are referred to as *shifts*. However during the two most recent reintroductions of H1N1 in 1977 and 2009, the new subtype established but did not replace the resident H3N2.

In addition to shifts in subtype, influenza A (and B) are subject to *drift* where point mutations in the HA and NA genes cause a gradual change in its antigenic properties with larger so-called cluster-jumps occurring every 4-5 years [26]. New drift-variants are referred to as *strains*. Influenza drift in combination with waning immunity means that reinfection with the same subtype typically occurs every 5-7 years [27]. From the view point of influenza evolution, influenza drift leads to a series of bottlenecks as successful drift-variants usually dominate global influenza circulation within a single season. Focusing on the HA-gene the resulting phylogenetic tree has a characteristic slender shape [28].

### Zooming in on the NS gene

In this study, we investigate a different aspect of influenza A evolution and look primarily at the NS gene, rather than the antigenically dominant HA or NA genes. We focus on the evolution of the NS gene since the devastating 1918 ‘Spanish flu’ pandemic, mainly viewing it through the lens of nucleotide distance maps. Our goal is to show how the the established origins of certain influenza strains are reflected in the evolution of the NS gene and how mathematically simple distance-based measures can complement traditional, tree-based phylogenetic inference in a highly transparent way.

That the *NS* of the NS gene stands for *nonstructural*, reflects a belief that the two proteins for which it codes (NS1 and NS2) are not part of the influenza virion, but only produced in infected cells. However, NS2 – also known as the Nuclear Export Protein (NEP) – has proven to be present in virions as well. It has long been known to play a role in facilitating the export of viral ribonucleoprotein complexes, and since it is not entirely nonstructural, the designation NEP is more fitting [29].

However, even with small amounts of NEP present in virions, the NS gene as a whole is believed to be under much less selective pressure than genes coding for prominent surface proteins, such as HA and NA, as these are primary targets for neutralizing antibodies of the host immune system [30], resulting in a strong pressure towards immune evasion by antigenic change. We focus on the NS gene precisely because it is only subject to weak selective pressures, and can thus be regarded as undergoing largely neutral genetic drift for long periods of time. Sudden deviations from neutral drift can then serve as indicators of e.g. changes happening in other genes (via the hitchhiking effect [31]).

### Nucleotide distance measures as evolutionary probes

Fundamentally, we use two different measures based on genotypic distance in our study of the evolution of the NS gene, each of which has its own strengths and provides distinct and complementary information.

The first method – the *rooted Hamming map* – shows the distances between a fixed reference sequence (the ‘root’) and all other sequences over time. This allows us to gauge the genetic (dis)similarity between the root and circulating sequences, as well as the rate of genetic divergence (i.e. the ‘molecular clock’). However, the ‘directionality’ of evolution is obscured in this view, since e.g. any two descendants of the root virus which are equally dissimilar to the root will appear ‘close’ in the resulting (time, distance) plot, despite having evolved independently since they split off from the root. Fundamentally, this owes to the fact that the dimensionality of the space of sequences is much larger than the one-dimensional space of distances onto which it is projected.

The second method – the *un*rooted Hamming distribution, as used in [32] – to some extent remedies this situation, since it measures the distribution of genotypic distances between simultaneously circulating viruses – a measure of sequence diversity. This allows us to ascertain the degree to which lineages have diverged from each other by independent genetic drift, or have evolved together (due to e.g. intermittent evolutionary bottle-necks).

It should be noted that the *un*rooted Hamming distribution requires rather large amounts of sequencing data to be feasible. The reason for this is that a sufficient number of independent samples are required within each time window, to build up a representative distribution for that time point. In practice, this means that the unrooted Hamming distibution can only be consistently generated using post-2008 data, due to the increased level of sequencing. In comparison, the unrooted Hamming distribution has been feasible for almost the entirety of the SARS-CoV-2 pandemic [32]. This highlights the value of extensive sequencing and how it enables new and qualitatively different analyses to be carried out.

While both methods compress *N* very high-dimensional data points (sequences) into a low-dimensional object (consisting either of *N* distances to a common reference or *N* (*N* + 1)*/*2 pairwise distances^1^), the two applications together give a rather detailed view of evolutionary speeds and within- and between-strain diversity. Viewed temporally, the methods therefore can also aid in uncovering evolutionary events such as genetic saltations, bottlenecks, recombinations and reassortments, at a glance.

### Long-term evolutionary trends

Applying the two sequence Hamming distance measures to the study of the NS gene over more than 100 years (1918-2023), we do indeed find that there has been a gradually evolving NS gene ‘backbone’ which has been present continuously within human influenza A (Figure 2), even during shifts from H1N1 (1918) to H2N2 (1957) and then to H3N2 (1968). As observed by [19] the 1977 reemergent H1N1-lineage follows the same backbone with a 20 year delay. However, the 2009 pandemic influenza H1N1 represents a sudden jump in genetic distance, compared to the previously circulating human H1N1. Interestingly, the descendants of the 2009pdm NS gene do not align themselves with the overall observed backbone, suggesting that it has either undergone neutral genetic drift at an accelerated rate, has been subject to selective pressures driving adaptive changes, or had a most recent common ancestor with the ‘backbone sequences’ some time prior to 1918.

**FIG. 1.**
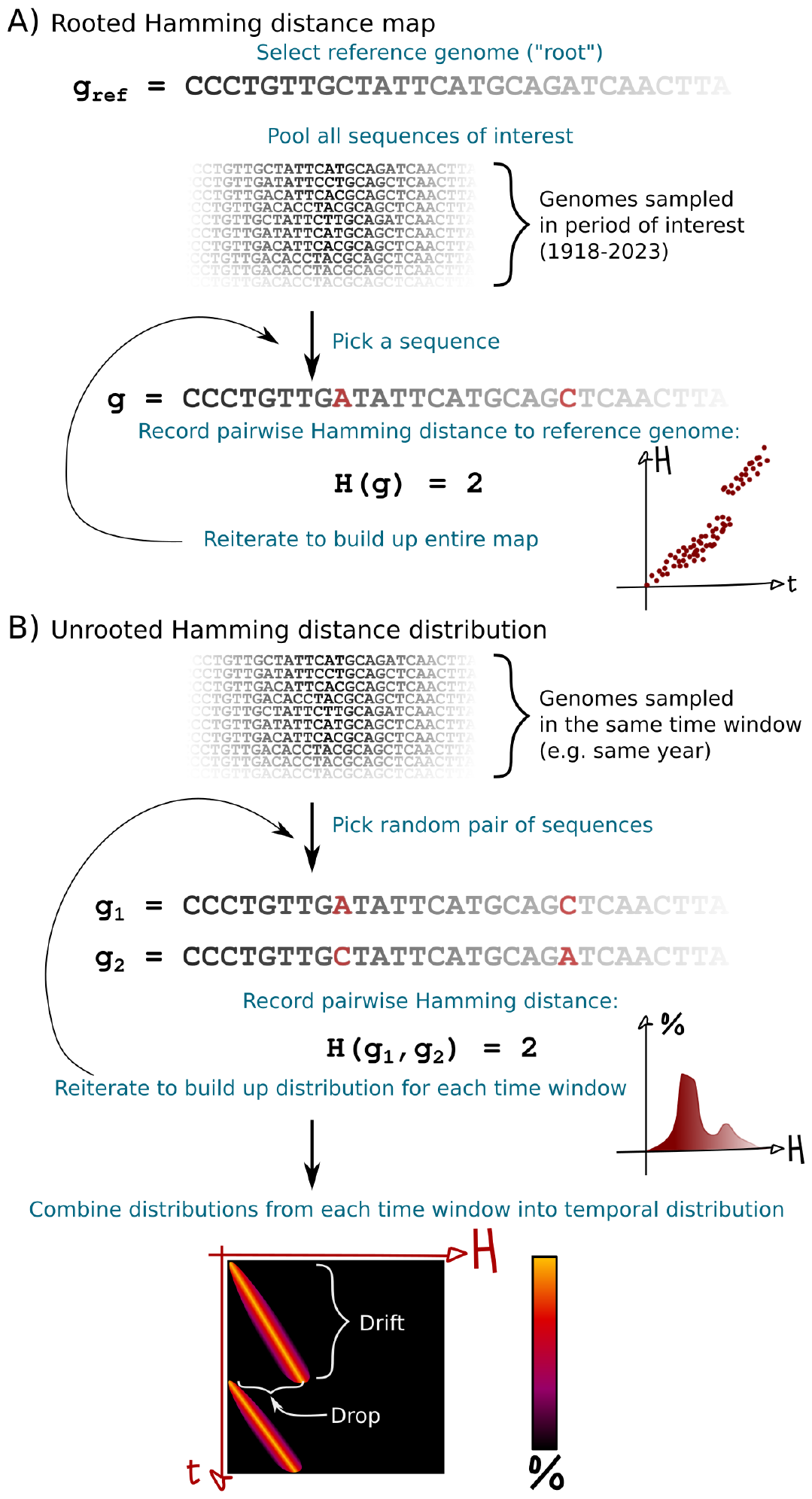
Data workflows involved in generating the temporal genotypic distance visualizations. **A)** The algorithm for generating a *rooted* Hamming distance map, used to visualize the similarity of a large set of sequences {*g*_*i*_}, sampled at different times, to some reference sequences *g*_ref_. **B)** Algorithm for generating an *unrooted* Hamming distance distribution, used to visualize temporal variations in sequence diversity among circulating (influenza) viruses.

**FIG. 2.**
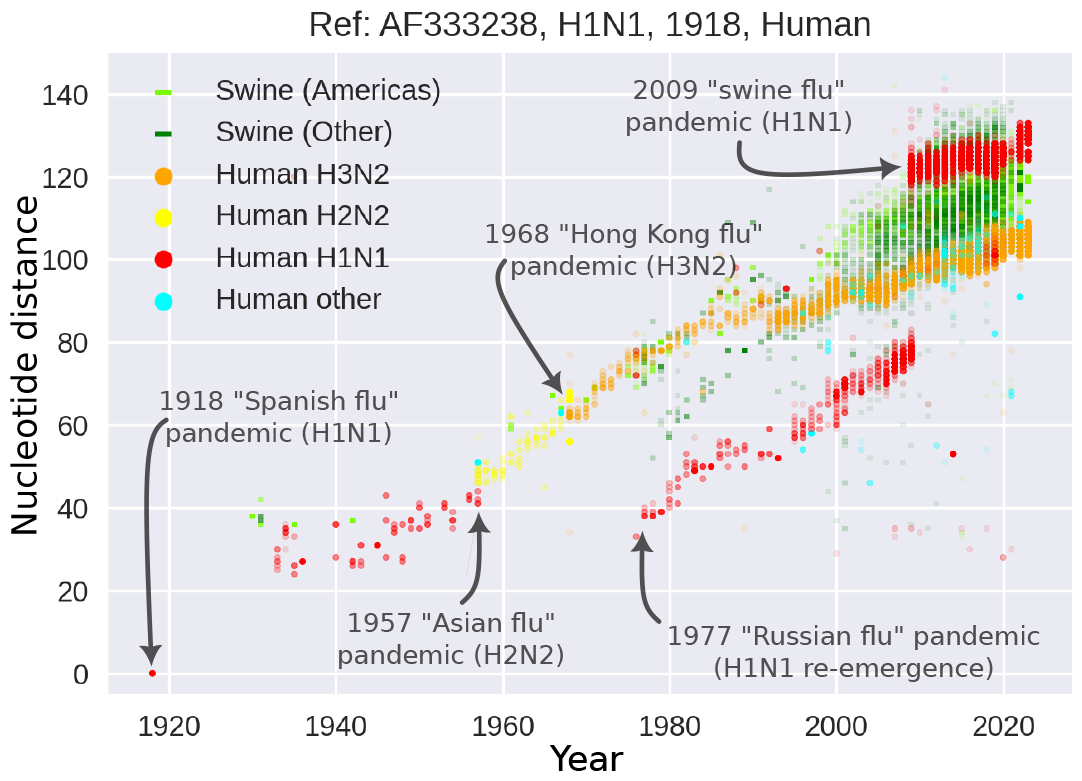
Nucleotide distances rooted in the 1918 Brevig Mission human-derived H1N1 sequence, GenBank accession number AF333238. A continuous ‘backbone’ is apparent, beginning with H1N1 subtype sequences (red) until 1957, then H2N2 (bright yellow) until 1968 followed by H3N2 (orange). In 1977, H1N1 reappears outside the backbone. This lineage is then supplanted by the 2009pdm H1N1 strain in 2009. Swine-derived sequences (green and black) are observed to diverge from the 1918 sequence over time as well, albeit with much larger variance.

In [19], the evolutionary rate of change of the NS gene segment of Influenza A H1N1 was investigated using a (rooted) nucleotide distance measure. As a reference (root) for distance calculations, they used a hypothetical sequence representing the origin of the best (maximally parsimonious) phylogenetic tree constructed from the sequences available to them. The oldest sequences used in [19] were from 1933 and 1934, respectively, while the newest was from 1985.

Their analysis demonstrated a linear increase in the total number of nucleotide sequences as a function of time, consistent with neutral drift at a well-defined molecular clock rate. We will use the analysis of [19] as a reference point, and discuss how their results hold up now that more than a century of influenza sequences are available. The largest deviations turn out to involve the 2009 H1N1 ‘Swine flu’ pandemic.

The current wisdom regarding the origin of the 2009 H1N1 pandemic influenza virus is complex, owing to how readily influenza viruses undergo reassortment. In a 2009 phylogenetic analysis, Smith et al. [33] estimate that the HA, NP, and NS genes as well as the polymerase genes (segments 1-3) emerged from an already triply reassortant virus circulating among swine in North America. This triple-reassortant itself contained genes from avian, human H3N2 and classical swine lineages. Most importantly for our purposes, the NS gene is believed to have been derived from classical swine lineages [33]. The NA and M genes, however, were estimated to originate in an Eurasian avian-like H1N1 lineage.

Garten et al. [34] go further and suggest that, ultimately, the ancestor of the NS gene of the 2009 H1N1 pandemic strain was transferred to pigs in 1918 and originated in birds. In this way, the NS gene of the 2009 H1N1 pandemic virus can be traced back, via a series of reassortment events, to the 1918 H1N1 pandemic virus where it is believed to have entered human and swine populations almost simultaneously [35]. Additionally, [28] studied the evolution of the HA1 domain of human H3N2 between 1984 and 1996 using raw nucleotide distance measures. However, their investigation was limited to a single subtype, and concerned a part of the genome which is under strong selection.

## II. METHODS AND MATERIALS

### NS gene sequence data

All the NS gene sequences used were obtained from the GenBank/NCBI Virus database [36]. Regardless of the host organism, our selection criteria were:

- Metadata must contain a date with at least the year specified.
- Minimum sequence length of 800 nucleotides
- Only influenza A is included.
- All samples collected up to the present day are included.

In the main part of the analysis, we include only samples obtained from either human or swine. This resulted in 61, 307 sequences. However, in Supplementary Figure 1 we include all host organisms for completeness, corresponding to 87, 796 sequences. It should be noted that humans, swine and avian species account for 97.3% of these sequences.

The code and data are available on GitHub [37].

### Temporal sequence Hamming distance distributions

We generate two different types of temporal Hamming distance visualization which we call ‘rooted’ maps and ‘unrooted’ distributions.

We stress that the two types of visualizations give entirely different but complementary information on the evolutionary dynamics as well as the population dynamics of the pathogen under scrutiny.

Below, we first describe the simple algorithms used to generate each type of temporal Hamming distance distribution and then discuss some of the qualitative properties of the distributions. The two methods are illustrated in Figure 1. Note that, as in [19], gaps in sequences do not contribute to the measured Hamming distances. Consequently, the Hamming distance represents the number of substitutions. In practice, this means that if there is a gap at a given position in one of the two sequences being compared, then that position is not counted toward the total Hamming distance between the two sequences. This choice was made to ensure that missing data in incomplete historical sequences would not dramatically skew the results.

#### The rooted Hamming map

The simplest visualization we use is the *rooted* Hamming map. The rooted Hamming map is defined in terms of some fixed *root* or *reference sequence g*_ref_. Once the root is chosen, the rooted Hamming map is constructed by computing the Hamming distance *H*(*g*) = *H*(*g, g*_ref_) for each sampled sequence *g*. Each sequence is then represented by a point of the form (*t*(*g*), *H*(*g*)), where *t*(*g*) is the collection time of the sample *g* and *H*(*g*) is the aforementioned Hamming distance. This workflow is illustrated in Figure 1A.

#### The unrooted temporal Hamming distribution

The second type of visualization that we employ is the unrooted temporal Hamming distribution, as developed in [32]. As the name implies, there is no fixed reference sequence in this case.

Instead, a moving time window of width Δ*t* is constructed, and all (or a large random subset) of the pairwise Hamming distances between sequences collected within each time window are computed.

Let

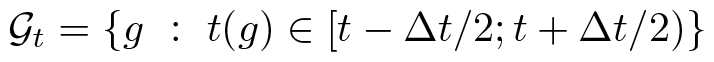

denote the set of all sequences *g* whose collection time *t*(*g*) lies within the time window of width Δ*t* centered on *t*. For each such set *𝒢*_*t*_, we then compute the distribution *P*_*t*_(*h*) of sequence Hamming distances *h* defined as

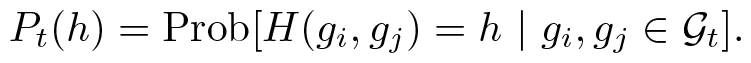

The object we have dubbed the unrooted temporal Hamming distribution is then simply the collection of these distributions in the form of

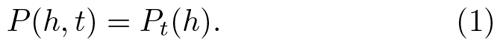

In practice, the time variable *t* is discretized (to e.g. days or years, depending on the sampling density and desired resolution) so that only a finite number of distributions *P*_*t*_(*h*) are computed.

As a consequence of equation 1, the normalization is such that 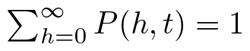 for any *t*.

The practical workflow involved in computing the unrooted temporal Hamming distribution is illustrated in Figure 1B.

#### Complementarity of the two metrics

Fundamentally, the rooted Hamming map yields clear visualizations of relatedness (or at least similarity) between a reference sequence and a large numbers of sequences at a glance. It can furthermore give indications of sudden, large changes in the genomes of circulating pathogens (as illustrated by e.g. the emergence of the 2009pdm H1N1 strain in this study).

The unrooted temporal Hamming distribution, on the other hand, gives access to information which is unobservable in the rooted Hamming map. For example, a sudden drop in diversity will be observable as an abrupt shift of the mass of the distribution *P* (*h, t*) towards smaller *h*-values, as shown near the bottom of Figure 1B. Thus, the unrooted temporal Hamming distribution can reveal population bottlenecks as well as variant transitions, even if the new variant is not genotypically very different from the previously circulating one. Such a transition would not be clearly visible from the simple rooted Hamming map. In fact, the unrooted temporal Hamming distribution can even yield information on the reproductive advantage of an emerging variant through the rate of growth of the (approximately binomial) peak corresponding to Hamming distance comparisons between genomes belonging to the new variant – i.e. the peak which appears at low Hamming distance at the invasion of a new strain.

The unrooted temporal Hamming distribution thus can discriminate between different scenarios that would look identical in the rooted Hamming map. One example of this is the evolution of 2009pdm-descendant H1N1 in swine and humans after 2009, which we discuss in this paper. The rooted Hamming map gives the impression that the virus evolved similarly in swine and humans after 2009, but the unrooted temporal Hamming distribution reveals very different patterns of viral diversity in the two populations, with implications for our understanding of the epidemiological dynamics of the two (see *Results* for details).

### Phylogenetic analysis

The timetree phylogeny in Figure 5 is created using BEAST2 [38]. The model used is described by the following elements:

- The site model consists of a categorical Gamma distribution of substitution rates between sites with 4 categories and an a priori unspecified shape parameter (to allow for rate heterogeneity).
- A Hasegawa-Kishino-Yano substitution model is used [39], with estimated overall substitution rate and value of *κ* (ratio between transition and transversion rates).
- A strict clock model is assumed.
- The prior distribution for the population size is taken as a constant-population Coalescent model.

Redoing the analysis assuming a Coalescent Bayesian Skyline model with 10 population sizes does not substantially affect the results. The mean time to the most recent common ancestor moves from 1890 to 1892 and the tree topology is unchanged.

## III. RESULTS

Nucleotide distance maps are low-dimensional projections of very high-dimensional data (the myriad ways in which two sequences can differ from each other). For this reason, viewing only a single such map rooted in one reference sequence can be deceptive: two different lineages which have diverged equally far from the root can appear close, despite having little similarity – they may, after all, have diverged in completely unrelated directions since branching off from their common ancestor.

However, if multiple such maps with different roots are combined, they can give a remarkably clear picture of the evolutionary relationships between multiple strains of the same pathogen.

### The 105 year influenza A backbone

The Hamming map of Figure 2 is rooted in the 1918 H1N1 NS sequence obtained at Brevig Mission, Alaska [2, 4]. It is worth noting that although sequences derived from swine are sparse before 1980, swine sequences from the 1930s are reasonably closely related to the 1918 strain, in terms of the NS gene (Figure 2). This observation is interesting when compared with the finding of [40] that mice that receive an A/Swine/Iowa/30-inactivated vaccine are completely protected against lethal challenge with recombinant influenza viruses possessing genes (at least HA and NA) from the 1918 influenza virus. These observations square well with the assessment of [34] that the NS gene of classical swine influenza was transferred to pigs around 1918. In the supplementary material, we include avian sequences as well – see Supplementary Figure 1. The supplementary plot includes the avian (chicken) sequences from 1902, Brescia, Italy, which are the oldest known influenza sequences [41]. While not conclusive, the supplementary plot suggests that the continuous NS gene ‘backbone’ observed from 1918 to 2009 can be extended backwards to 1902. In Figure 2, the 20^th^ and 21^st^ century pandemics associated with the dominance of a new subtype have been indicated.

In Figure 3, we have collected several snapshots in the form of Hamming distance maps rooted in nine different reference sequences. This allows us to build up a natural history of human influenza A for the last 105 years and to look at the events indicated in Figure 2 in more detail.

**FIG. 3.**
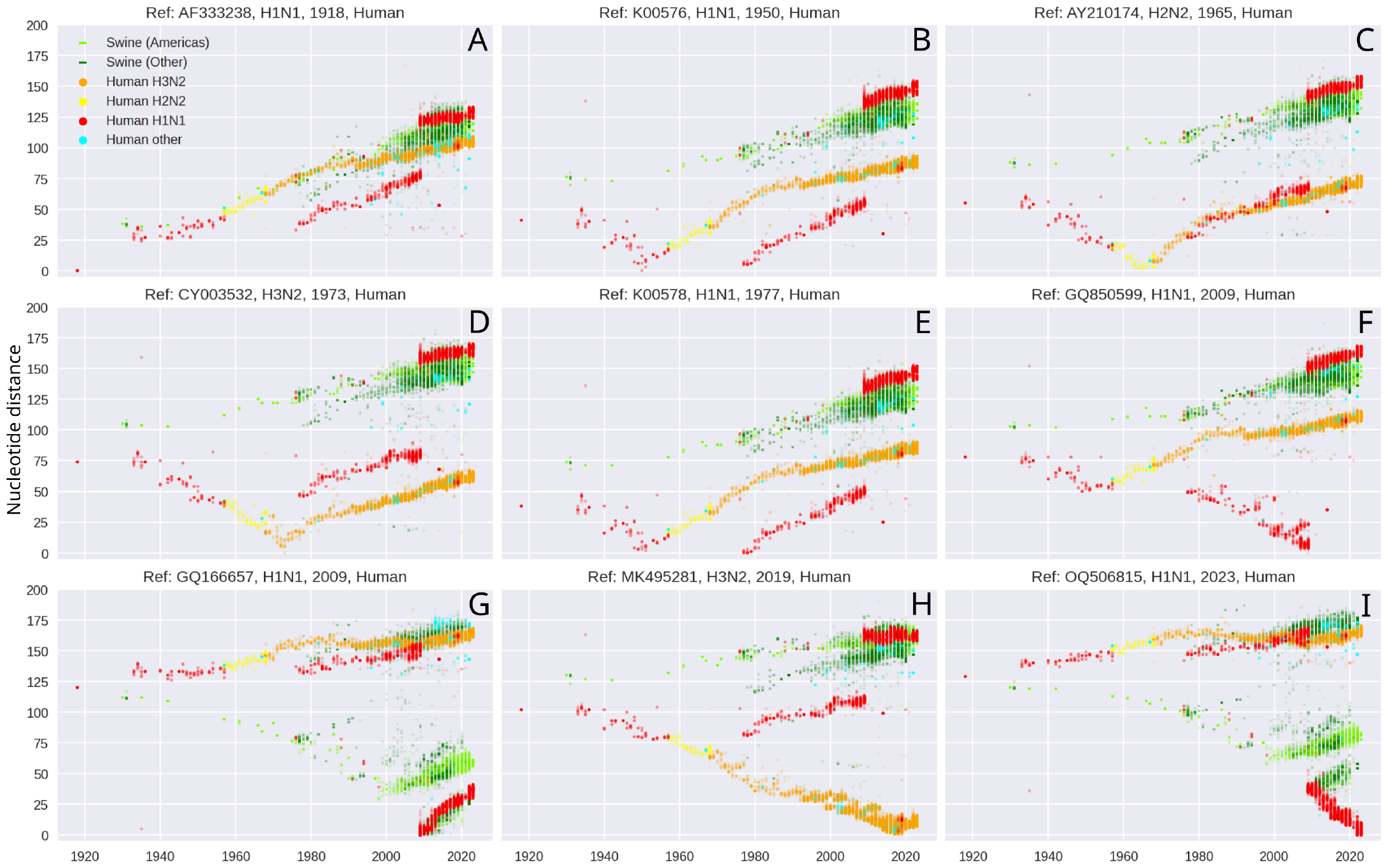
105 years of influenza NS gene evolution in humans and swine. In each panel, we root the nucleotide distance analysis in a different reference sequence. **A)** Rooted in the 1918 Brevig Mission H1N1 sequence. **B)** Rooted in a 1950 US H1N1 sequence. Note the similarity to 1977 H1N1 sequences. **C)** Rooted in a 1965 US H2N2 sequence. **D)** Rooted in a 1973 Hong Kong H3N2 sequence. **E)** Rooted in a 1977 USSR H1N1 sequence. Note the similarity to H1N1 circulating in the early 1950s. **F)** Rooted in a 2009 Chinese H1N1 sequence belonging to the 1977 clade. **G)** Rooted in a 2009 UK H1N1 sequence, belonging to the 2009pdm strain. **H)** Rooted in a 2019 Russian H3N2 sequence, retrospectively displaying the unbroken genetic ‘backbone’. **I)** Rooted in a 2023 US H1N1 sequence of the 2009pdm clade.

The main observation from panel A is now well-known (from Figure 2): the 1918 pandemic strain is seemingly an ancestor of the bulk of later strains, since those fall more or less on a straight line. The notable exception to the above pattern is the H1N1 lineage that begins in 1977. Panels B and E together show that the 1977 lineage is almost identical to sequences sampled in the early 1950s, as noted by [19, 42].

Panel C shows that the NS gene of the H2N2 virus which emerged in Southern China in 1957 (‘the Asian flue’) and caused a deadly pandemic is a direct descendant of the H1N1 lineage that preceded it. These ‘retrospective’ plots are important (i.e. looking back from 1957), because they allow us to distinguish between two scenarios which the ‘prospective’ plot of panel A cannot delineate:

*Scenario A)* The NS gene of the 1957 H2N2 strain is a direct descendant of the H1N1 strain that circulated just prior, which in turn descends from the 1918 H1N1 strain. *Scenario B)* The NS genes of the 1957 H2N2 strain and the H1N1 strain that circulated just before it have diverged equally far from the 1918 H1N1 strain, but are not closely related to each other.

Figure 3C shows that the former interpretation is the correct one.

Similarly to the analysis for H2N2 above, panel D shows that the H3N2 ‘Hong Kong’ pandemic strain which emerged in 1968 inherited its NS gene from the previously circulating H2N2.

In panel F, we root the analysis in a 2009 H1N1 strain which is *not* of the 2009pdm-type. Viewed retrospectively (i.e. looking at sequences prior to 2009), the plot shows that this lineage, which would soon be out-competed by the emerging 2009pdm strain, can be traced back to the 1977 re-emergence of H1N1 (and thus ultimately back to the 1918 strain).

Conversely, in panel G, we root the analysis in a 2009pdm sequence. It is clear that the NS gene of H1N1 2009pdm and the previously circulating human H1N1 are only very distantly related, differing by around 150 nucleotides.

In panel H, we wrap up the story as far as the NS gene of human H3N2 is concerned, by rooting in a relatively recent (2019) H3N2 sequence. This plot reveals that the H1N1-H2N2-H3N2 backbone has long diverged from other contemporary strains and by now exists in a rather isolated region of the sequence space.

Lastly, panel I is another ‘retrospective’ plot in the sense that it allows us to disambiguate a feature of Figure 3G. While it was not clear from panel G whether the H1N1 2009pdm strain and the closely related swine-derived lineages had evolved in the same ‘direction’ since 2009 (suggesting back-and-forth transmission), panel I shows that this is not the case. Rather, the H1N1 2009pdm-descendants circulating in humans have diverged from their closest co-circulating swine-derived counterparts at a steady rate since 2008 or 2009. We look more closely at this situation in Figure 4 with the help of unrooted Hamming distributions.

**FIG. 4.**
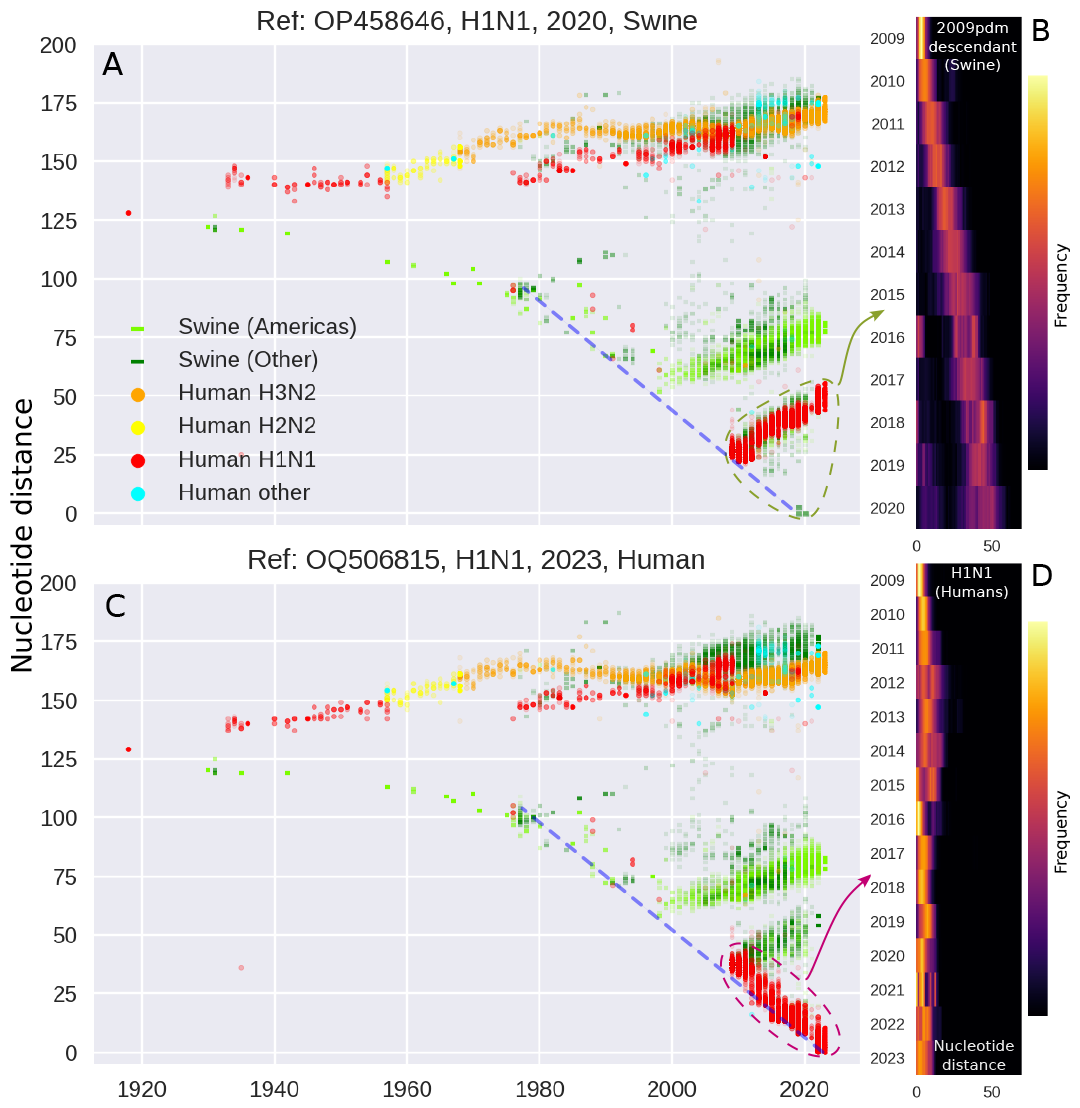
**A)** NS gene nucleotide distance map rooted in a 2020 swine-derived H1N1 sequence. **B)** Unrooted (pairwise) Hamming distance distribution for swine sequences sampled from 2009-2020. Note the approximately linear increase in the characteristic distance indicating growing diversity with-out severe bottlenecks. **C)** NS gene nucleotide distance map rooted in a 2023 Human H1N1 sequence. **D)** Unrooted (pair-wise) Hamming distance distribution for human sequences from 2009 onward. Note that – in contrast to panel B – a virtually constant low level of diversity is observed, indicating yearly bottle-necks.

**FIG. 5.**
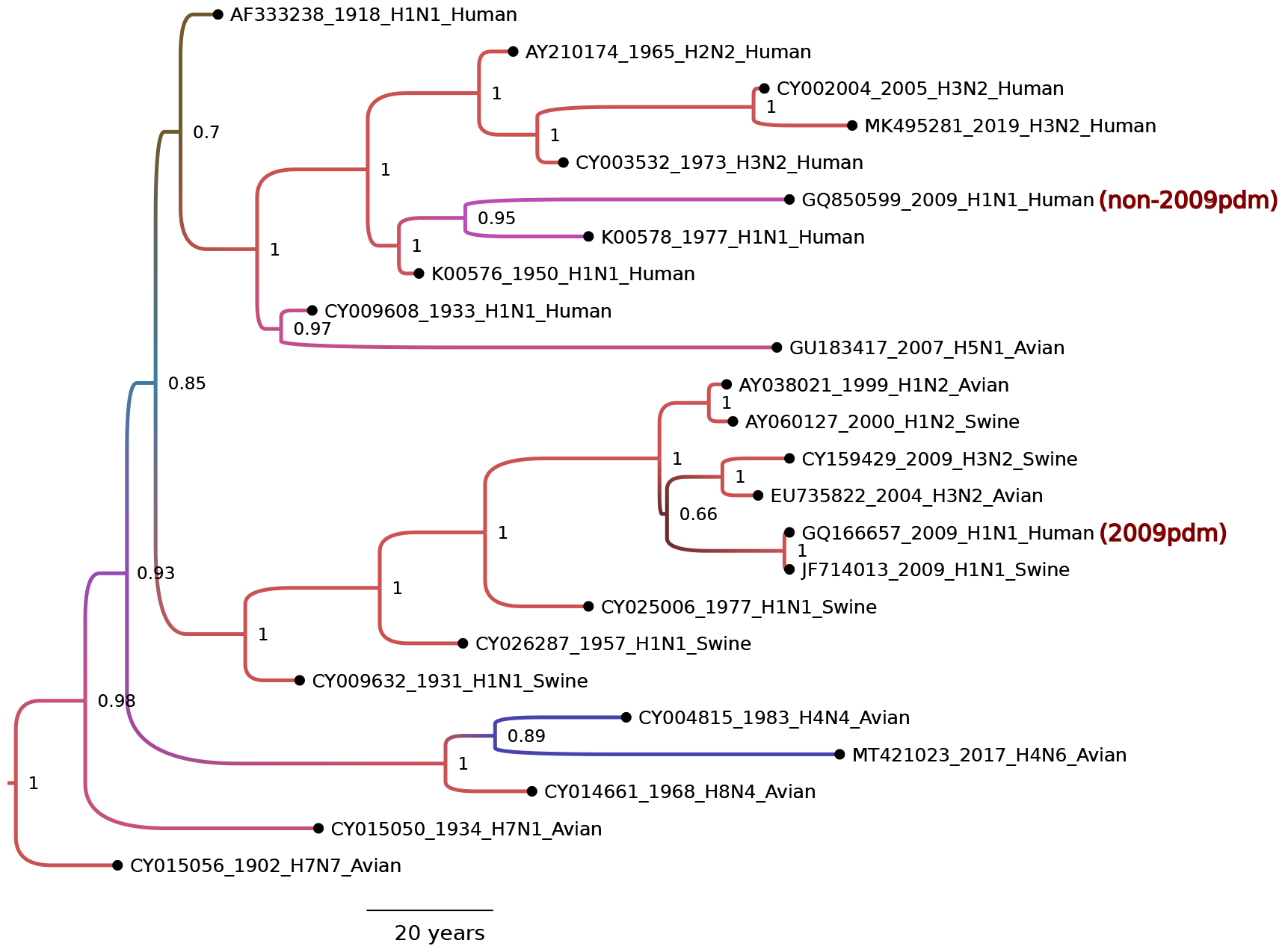
Maximum-likelihood phylogeny (timetree) for the NS gene in human H1N1, H2N2 and H3N2. Avian and swine-derived sequences (regardless of influenza A subtype) with high similarity to human sequences were included to probe the origin of the NS gene of the 2009pdm strain in humans. Colored by posterior (lowest value shown: 0.66, highest: 1.0). Each sequence is labeled by its accession number, year, subtype and host. The analysis places the most recent common ancestor of the sampled NS genes (including the 1918 Human H1N1 and the 1902 avian H7N7) around 1890 (95% highest probability density (HPD) interval: [1882,1898]).

### The 2009 swine influenza viewed through the (un)rooted Hamming distribution

Figure 4A is rooted in a representative reference sequence of swine-host 2009pdm-descendant H1N1 NS genes. The figure demonstrates that a straight line can be drawn connecting a punctuated backbone of swine isolates going back to the late 1970s to the 2009pdm sequences existing in humans as well as swine, and all the way to present-day swine isolates. In Panel B, we show the pairwise (“unrooted”) nucleotide distance distribution based on swine-derived sequences located in the lower right-hand corner of panel A (the green points enclosed by a green dashed line), i.e. sequences from 2009 onward which are likely descendants of the original 2009 strain. We observe that the pairwise Hamming distance increases linearly from almost zero in 2009 to a typical distance of 45-50 in 2020. This indicates that the circulating 2009pdm-descendant lineages in swine have continuously diverged from each other during this time, without suffering any severe bottlenecks (which would show up as sudden drops in diversity, i.e. drops in the pairwise Hamming distance).

Comparing with Figure 4C, which is rooted in a 2023 human pdm2009-descendant H1N1 sequence, the initial observation is the same: a straight line can be drawn back to the late 1970s, making contact with the original 2009pdm strain and several swine sequences obtained between 1977 and the late 1990s/early 2000s. However, the year-by-year *un*rooted Hamming distribution based on human-derived 2009pdm-descendant H1N1 sequences (the red points enclosed by a dashed red line in panel C) shown in Figure 4D is remarkably dissimilar to the one for swine shown in Figure 4B. The human-host Hamming distribution shows signs of repeated bottlenecks, with the typical pairwise Hamming distance barely rising above 5-10 nucleotides, indicating that genetic diversity remains low throughout the period. This is in stark contrast to the swine-host Hamming distribution which shows linearly increasing diversity, indicative of largely neutral genetic drift free of any severe population bottle-necks.

A likely explanation for the limited standing diversity of 2009pdm-descendant H1N1 in humans is the seasonal nature of the disease and the selective pressure arising from population immunity. Due to the shorter lifespan and resulting high population turnover rate among farmed swine, population immunity to influenza is limited in these populations. Correspondingly, it has been noted that influenza in modern farmed swine does not have the same strong seasonal bottlenecks [43, 44] as in the human population. This lack of strong seasonality in swine likely owes to the high concentrations and rapid population turnover of modern farming practices.

The limited standing diversity of the human-host 2009pdm-descendant virus population also explains another feature which differs between Figure 4A and 4C. In Figure 4C, a thick line of intermediate sequences leads from the original 2009pdm strain (at distance ≈ 35) down to the root sequence. In the corresponding swine-host plot of 4A, no such string of intermediate sequences can be seen. However, this is only natural given the distributions of panels B and D. With limited standing diversity, any sampled sequence in the intermediate years will be fairly closely related to a contemporary ancestor of the 2020 root sequence, giving rise to the clear string of intermediate sequences. In contrast, unhindered genetic drift with an absence of bottlenecks leads to every lineage diverging from every other at a steady pace. This means that sequences belonging to different transmission chains have likely continually diverged from each other since around the time of the last severe bottleneck, i.e. since 2009. It follows that sampling in the swine population at a given time yields a very small probability of having sampled from the same transmission chain in earlier years (post-2009) simply due to the large volume of the sequence space.

A similar line of reasoning may explain why no close ancestors to the 2009pdm strain can be observed in swine in the period between 2000 and 2008. Unfortunately, the level of sequencing is not high enough that we can establish an unrooted Hamming distribution for the relevant section of Figure 4C for those years. However, assuming that influenza in swine also had only limited seasonality (and thus few severe bottlenecks) during that period, the argument above explains why we would not expect to detect a close ancestor among the sampled sequences.

### Phylogenetic analysis

The timetree phylogeny in Figure 5 (created using BEAST2 [38], see Methods and Materials for details) corroborates two of the observations already suggested by the Hamming maps of Figure 4: 1) in terms of the NS gene, classical swine flu and the human lineage which gave rise to the H1N1-H2N2-H3N2 backbone already diverged around the time of – or slightly before – the 1918 ‘Spanish flu’ pandemic. 2) The NS gene of the 2009 H1N1 pandemic in humans seems to originate from classical swine flu, as found by [33].

In the phylogeny, we have included several historic avian influenza sequences as well, including one from a collection dating back to 1902, Brescia, Italy – the oldest known influenza sequences [41]. As can be seen in the Supplementary Figure 1 as well as the phylogeny, there are some late-1990s and early-2000s avian sequences (of various subtypes) which are remarkably closely related to 2009pdm. Already in 2002, Suarez et al. [45] studied an H1N2 influenza virus isolated from turkeys and found it to be a complex reassortant virus with a mix of swine-, human-, and avian-origin influenza genes. This sequence has accession number AY038021 and can be found in the phylogeny of Figure 5, intermingled with human and swine sequences. Along the same lines, Choi et al. [46] study two 2003 isolates of H3N2 from turkeys and find them to have 97% to 98% sequence identity to swine H3N2 viruses, demonstrating interspecies transmission between pigs and turkeys. This, combined with the intermingling of viruses isolated from swine, humans and avian species in the region of the phylogeny where the 2009pdm strain arises, underscores the significant uncertainties involved in establishing the precise path of a new viral strain into the human population.

## IV. DISCUSSION

Buonagurio et al. named their 1986 paper [19] *“Evolution of human influenza A viruses over 50 years: Rapid, uniform rate of change in NS gene”*. As the title suggests, one of their main findings was that the NS gene had essentially diverged at a steady pace from around 1933 – the earliest sequence available to them – and up to the then-present day. This observation of course came with the caveat that the emergence of the 1977 H1N1 represented an apparent departure from the temporal pattern, representing a sudden drop in nucleotide distance to the reference sequence. However, realizing that the 1977 sequence is all-but identical to early 1950s H1N1 (likely the result of laboratory escape [42, 47–49]), it may be considered a continuation of the existing backbone, and indeed falls directly onto the existing line when translated horizontally, backwards in time, by approximately 27 years.

Along the same lines, the authors of [19] report that “there was evidence that the NS gene was not exchanged [50] during the reassortment of influenza viruses leading to new A virus subtypes [51]”. Since then, the pattern has evidently changed. We observe an abrupt change in the circulating H1N1 NS genes at the emergence of the 2009pdm strain, and this sudden shift is clearly visible in the nucleotide distance plots of Figure 4. Correspondingly, the phylogeny of Figure 5 shows that the NS gene of the H1N1 2009pdm circulating in humans originated with classical swine influenza, rather than variants previously circulating among humans.

The 2009 H1N1 pandemic was ‘weird’ in other respects as well. In 2001, Potter wrote “[…] a pandemic is caused by a new influenza virus A subtype, the HA of which is not related to that of influenza viruses circulating immediately before the outbreak, and could not have arisen from those viruses by mutation” [23], echoing Webster and Laver (1972) [52] who wrote “The other kind of antigenic variation (major antigenic shifts) involves sudden and complete changes in one or both of the surface antigens so that ‘new’ viruses arise to which the population has litle or no immunity and it is these viruses that are causes of the major pandemics of influenza”. However, Influenza A H1N1 was circulating widely in the human population immediately prior to the emergence of the H1N1 2009 pandemic strain.

One possible response to this observation would be to point out that the 2009 ‘swine flu’ pandemic was a relatively feeble one, as pandemics come. The mortality estimates were comparable to a seasonal influenza [53, 54], although the age distribution of mortality was skewed towards younger individuals (*<* 65 years) so that many more years of life were lost [53]. Indeed, antigenic similarity between the 2009pdm strain and previously circulating H1N1 strains may have have played a role in explaining the age-related immunity to the 2009 strain [55]. As such, the 2009 ‘Swine flu’ represents somewhat of a middle-ground between seasonal recurrence of similar influenza strains and the novelty of newly-emerged subtypes seen in previous influenza pandemics. See also Supplementary Figure 2, which shows that the nucleotide distances between the HA gene of the 2009pdm strain and previously circulating human H1N1 strains are much lower than those between the 2009pdm strain and e.g. previously circulating H3N2 or H2N2 strains. Thus, in 1977 as well as in 2009 the new strain was closely related to strains that had circulated only a few decades earlier – this may explain why the subtype established without causing a true shift where one would expect the resident subtype to be replaced (i.e. H3N2 continued to circulate among humans after 2009).

In the *Results*, we remarked that the Hamming distance analysis seemed compatible with the continuous NS gene ‘backbone’ observed from 1918 to 2009 extending backwards to at least 1902 (the earliest known influenza sample, which is of avian origin). Interestingly, the phylogenetic analysis behind Figure 5 results in a most recent common ancestor for all the included NS genes around 1890 (95% highest probability density (HPD) interval: [1882,1898]). This suggests an influenza spillover event or severe viral population bottleneck occurring around that time. This is notable because of the 1889-1890 ‘Russian flu’ pandemic, whose origins are still contested. While traditionally assumed to be an influenza pandemic, some scholars have suggested that it may be a coronavirus, most likely OC43 [56–58]. While certainly not conclusive, our analysis of the evolution of the NS gene is compatible with the hypothesis that a significant *influenza* spillover event happened around that time, coinciding with the 1889-1890 pandemic. However, other explanations for a coalescence date around 1890 are conceivable, including external factors leading to a major bottleneck in the viral population at the time. If the OC43 hypothesis turns out to be the most likely, then transient, pathogen-transcending (innate) immunity may be investigated as a possible explanation for an influenza virus bottleneck occurring around the same time [24, 59].

## V. CONCLUSION

Spurred on by remarks made by Freddy B. Christiansen – in turn inspired by [19] – we have followed the evolution of the human influenza A NS gene from 1918 to the present day, with the use of raw nucleotide distance measurements. Two decades have passed since Freddy suggested the analysis, and much has happened in terms of influenza evolution. The 2009 H1N1 pandemic, with its complex origins, was an especially dramatic departure from the linear trend hitherto observed (in e.g. [19]). The nearest origin of the 2009 pandemic proved clearly visible using the rooted Hamming map, and with the help of the unrooted Hamming distribution, we could shed light on some likely eco-evolutionary explanations for the lack of any clear (sampled) ancestor to the 2009pdm strain in the decade leading up to the 2009 pandemic.

The analysis using Hamming distance-based measures gives a simple and intuitive tool to study pathogen evolution that is very much in the spirit of Freddy B. Christiansen’s work. While the nucleotide distance maps can only complement more elaborate Bayesian phylogenetic inference methods, we believe that their model-free and directly interpretable nature holds great value.

## VI. ACKNOWLEDGEMENTS

BFN, CB and VA acknowledge financial support from the Danish National Research Foundation (grant no. DNRF170) and from the Carlsberg Foundation under its Semper Ardens programme (grant no. CF20-0046). BTG acknowledges financial support from the Flu Lab, The Princeton Catalysis Initiative, and the Schmidt DataX Fund at Princeton University made possible through a major gift from the Schmidt Futures Foundation. BFN furthermore acknowledges financial support in the form of a Carlsberg Foundation Internationalisation Fellowship (#CF23-0173).

## SUPPLEMENTARY MATERIALS

**Supplementary Figure 1.**
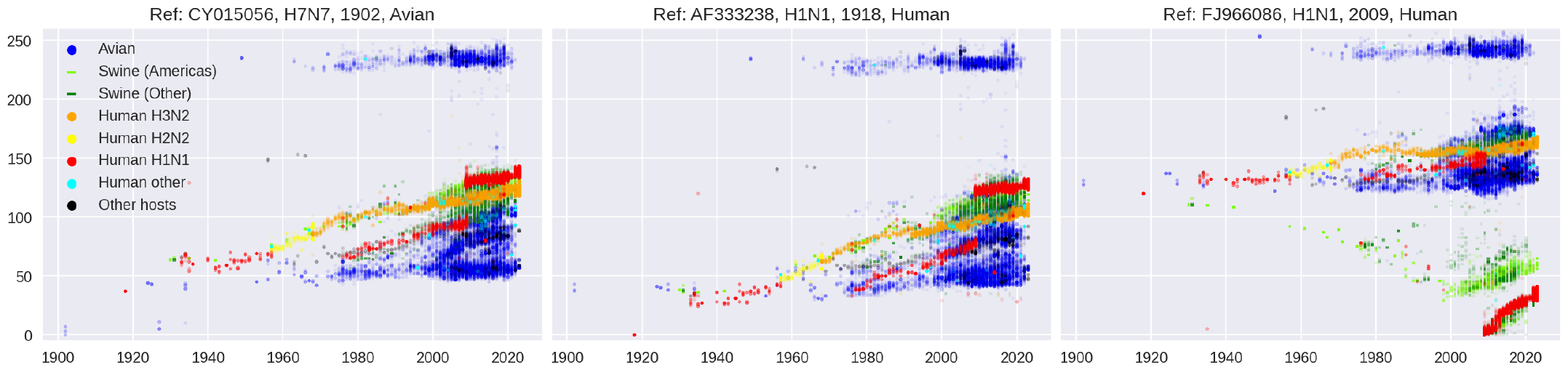
Rooted Hamming map of the NS gene, including all host animals. Here, GenBank sequences derived from all host organisms are included. Humans, swine and avian species comprise 97.3% of these 87,796 NS gene sequences.

**Supplementary Figure 2.**
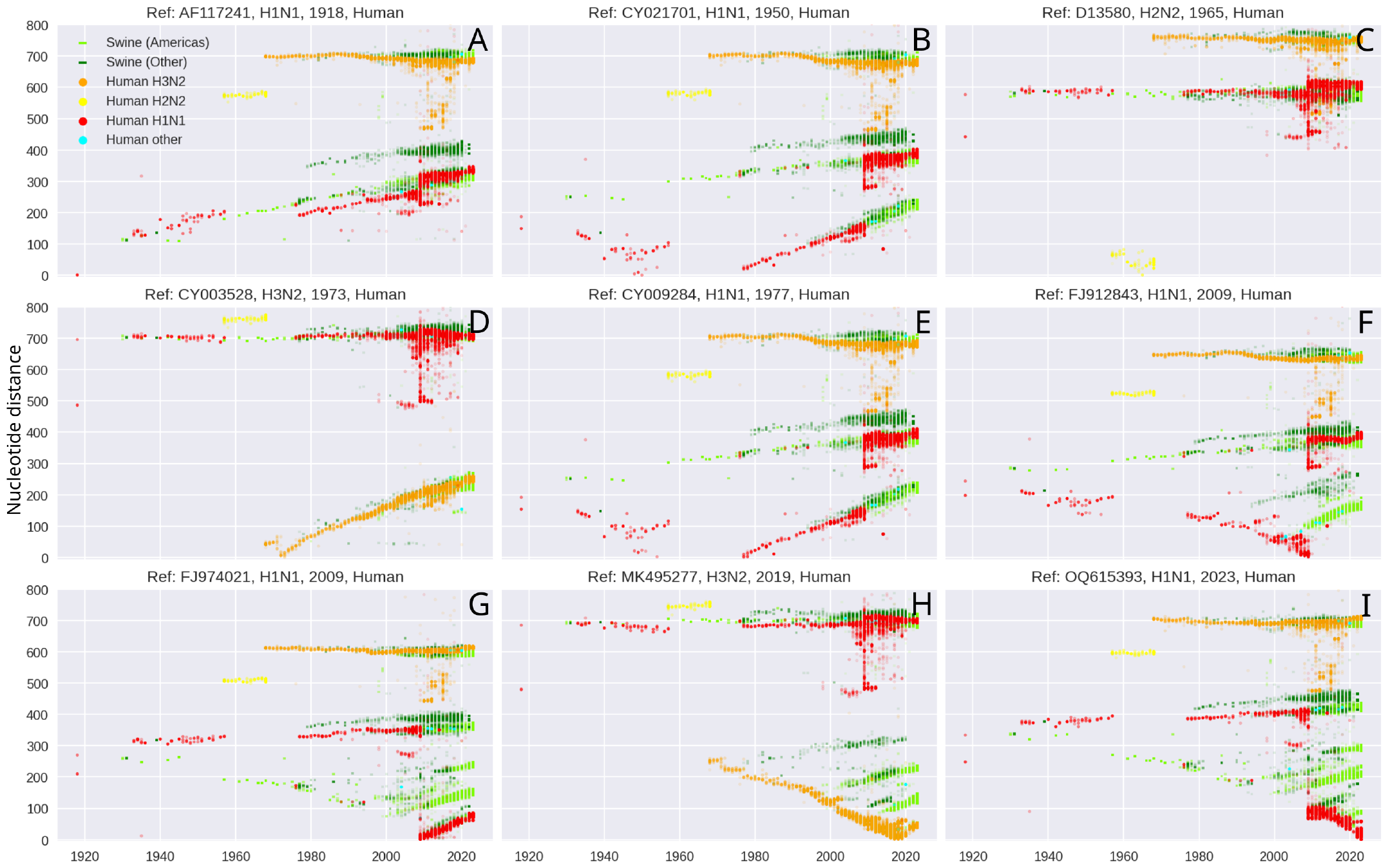
105 years of influenza HA (*Hemagglutinin*) gene evolution in humans and swine.

## Footnotes

1 It could also be argued that the two methods really produce *N* − 1 and *N* (*N* − 1)*/*2 distinct distances, if the (necessarily vanishing) distance between a sequence and itself is not counted.

